# Combination of photoactivation with lattice light-sheet imaging reveals untemplated, lamellar ruffle generation by PA-Rac1

**DOI:** 10.1101/2020.09.01.276824

**Authors:** Finian Leyden, Sanjeev Uthishtran, U K Moorthi, H M York, A Patil, H Gandhi, EP Petrov, S Arumugam

## Abstract

Membrane protrusions that occur on the dorsal surface of a cell are an excellent experimental system to study actin machinery at work in a living cell. Small GTPase Rac1 controls the membrane protrusions that form and encapsulate extracellular volumes to perform pinocytic or phagocytic functions. Here, capitalizing on rapid volumetric imaging capabilities of lattice light-sheet microscopy (LLSM), we describe optogenetic approaches using photoactivable Rac1 (PA-Rac1) for controlled ruffle generation. We demonstrate that PA-Rac1 activation needs to be continuous, suggesting a threshold local concentration for sustained actin polymerization leading to ruffling. We show that Rac1 activation leads to actin assembly at the dorsal surface of the cell membrane that result in sheet-like protrusion formation without any requirement of a template. Further, this approach can be used to study the complex morpho-dynamics of the protrusions or to investigate specific proteins that may be enriched in the ruffles. Deactivating PA-Rac1 leads to complex contractile processes resulting in formation of macropinosomes. Using multicolour imaging in combination with these approaches, we find that Myo1e specifically is enriched in the ruffles.

## INTRODUCTION

Cells orchestrate several dynamic plasma membrane structures to meet various functional demands. Macropinocytosis is one such fundamental cellular process, involving complex membrane dynamics driven by the actin cytoskeletal network enclosed within the membrane protrusions (1-3). A variety of molecular modules, including small GTPases, cytoskeletal nucleators and remodelling proteins, in conjunction with the actin cytoskeleton, comprise the ‘lamella-machinery’ required to generate, sustain and complete the ruffling resulting in the formation of a macropinosome or evince other physiological processes (4-7). Several small GTPases including Rac, Rab5 and Rab34 are implicated in the formation of ruffles (8-10).

The formation of dorsal ruffles or any elaborate membrane protrusion structure on the dorsal surface of the cell is a sophisticated multifaceted event that involves complex spatio-temporal activity of protein machineries (7). It is poorly understood how the ‘lamella-machinery’ utilizes biochemical and biophysical interplay to execute the complex membrane ruffling and closure required to form macropinosomes. Several studies have described the role of Rac1 in the formation of membrane ruffles and macropinosomes. Expression of the inactive mutant of Rac1 (Rac1T17N) inhibits macropinosomes, whereas the dominant active mutant (Rac1G12V) stimulates macropinosomes (11, 12). Indirect aggravation of Rac1 activity through constitutively active p21-acivated kinase induces macropinosome formation (13). The elucidation of activities regulating Rac1 was furthered by photoactivable Rac1 (PA-Rac1), a light-sensitive version of Rac1, created by fusing dominant negative Rac1 with the light oxygen voltage (LOV) domain and the carboxy-terminal helical extension of phototropin 1(14, 15). PA-Rac1 was used to decipher the role of activation and deactivation of Rac1. PA-Rac1 expressed in Raw264 macrophages induced membrane ruffling and macropinocytic cup formation (16). They further demonstrate that deactivation of Rac1 is crucial for the progress of pinocytic cup closure. In line with this model of Rac1 switching, in the structurally analogous process of phagocytosis, Ikeda et al. show that Rac1 switching at the correct spatio-temporal instance is necessary for phagosome formation (17). In general, the phagosome or macropinosome formation can be simplified into a two-step process – the first step of extending the membrane protrusion three-dimensionally, and the second step of engulfment/wrapping around leading to closure. All the previous evidence of Rac1 being constitutively active leading to defects in the closure of macropinocytic or phagocytic cups suggests that Rac1 switching is necessary to switch from the growth phase to engulfment and closure phase. Recently, Fritz-Laylin et al. used LLSM to image neutrophil-like HL-60 cells and found two important aspects of lamellar protrusions that formed in the absence of any substrate interactions: 1. Free standing lamellae combine to give rise to complex geometries that are essentially lamellar, and 2. Lamellar protrusions do not require any ‘surface template’ (18).

Investigating the morphodynamics of membrane protrusions formed on the dorsal surface of the cell is challenging. Despite the accumulation of much data on the phenomenology of membrane lamellipodial and ruffle formation, it has been difficult to image the membrane ruffles formed at the dorsal surface of the cell. All these characteristics demand imaging techniques to be capable of acquiring rapid volumetric imaging that minimize photo-bleaching effects. In the present report, we set out to use lattice light-sheet microscopy (LLSM) in combination with optogenetics to investigate PA-Rac1−mediated membrane protrusions on the dorsal surface and investigate the spatio-temporal events leading to Rac1 activation cycling-mediated macropinosome formation. An optogenetic approach, in which a photo-activatable chimeric construct alters to near-native functionality upon activation by light, is valuable to study such protein-based switches in complex protein machinery systems, and in combination with lattice light sheet that enables fast, volumetric live cell imaging, it becomes a powerful tool to investigate the processes on the dorsal membrane of the cell.

Using this experimental system, we demonstrate that:

1. Continuous activation of PA-Rac1 is required for effective ruffle formation on dorsal surface; sequential photoactivation re-initiates polymerization induced growth of pre-formed ruffles.
2. PA-Rac1-based ruffles are intrinsically sheet-like and can be generated spontaneously on the dorsal surface or can lift-off from lamellipodia attached to substrate.
3. Optogenetic stimulation of PA-Rac1 allows characterizing various morphodynamical features as well as ruffle associated proteins.

## METHODS

### Cell Culture

RPE1 (ATCC) cells were incubated at 37°C in 5% CO_2_ in high glucose Dulbecco’s modified eagle media (DMEM) (Life Technologies), supplemented with 10% fetal bovine serum (FBS) and 1% penicillin and streptomycin (Life Technologies). Cells were seeded at a density of 200,000 per well in a 6-well plate containing 5 mm glass coverslips. Phorbol 12-myristate 13-acetate (PMA) activation was performed by adding PMA to the imaging chamber at 100 nM.

### Plasmids and transfection

Cells were transfected with pTriEx-mCherry-PA-Rac1, a gift from Klaus Hahn (Addgene plasmid # 22027) (14), pGFP-EB1, a gift from Lynne Cassimeris (Addgene plasmid # 17234) (19), Myo1E-pmAppleC1, a gift from Christien Merrifield (Addgene plasmid # 27698) (20), Myosin-IIA-GFP, a gift from Matthew Krummel (Addgene plasmid # 38297) (21), EGFPC1-hMyoX, a gift from Emanuel Strehler (Addgene plasmid # 47608) (22), pTriEx-mVenus-PA-Rac1, a gift from Klaus Hahn (Addgene plasmid # 22007) (14).

Cells were transfected with 1 µg DNA (plasmid of interest – 0.2 µg, blank DNA – 0.8 µg) for single protein expression or (plasmids of interest – 0.2 µg + 0.2 µg, blank DNA – 0.6 µg) using lipofectamine 3000 (Thermo Fisher Scientific).

### Microscopy

Cells were imaged using a lattice light-sheet microscope (3i, Denver, Colarado, USA). Excitation was achieved using 488, 560 and 640 nm diode lasers (MPB communications) at 1-5% AOTF transmittance through an excitation objective (Special Optics 28.6× 0.7 NA 3.74-mm immersion lens) and was detected via a Nikon CFI Apo LWD 25× 1.1 NA water immersion lens with a 2.5x tube lens. Live cells were imaged in 8 mL of 37°C-heated DMEM and were acquired with 2x Hamamatsu Orca Flash 4.0 V2 sCMOS cameras. For SLM-based optogenetic excitation, 445 nm with full or quarter mask patterns was included in the sequence with 100% laser power. For wide-field excitation, a 1W 445 nm was attenuated with neutral density filters, expanded and focused on the back focal plane using a 700 mm focal length laser. Laser powers were modulated using a neutral density filter wheel as measured at the back focal plane of the collection objective. Lasers were blocked physically using a beam blocker (Thor labs) manually by keeping time using number of volumes acquired and recorded. Images were acquired using 3i Slidebook and deskewed using custom codes and deconvolved using Richardson-Lucy algorithm.

### Image analysis and segmentation

To quantify ruffle lengths, maximum intensity projection images were generated and analysed using FIJI. The images, after thresholding, were skeletonized and parameters were extracted using he plugin ‘analyse skeleton’ (23). To measure the ruffle thickness, profiles across the cross section of ruffles from deconvolved single planes of the volumetric stack were measured. To measure the contour lengths, SyGlass VR was used (24). Images were surface rendered, and the contour lengths were drawn manually. The collated data was exported, and end-to-end distance was extracted from the first and last coordinates of each ruffle. The dependences of the squared end-to-end dstance on the contour length were fit with a 2D worm-like chain model (25). Similarly, heights were measured manually along the length using the surface rendered boundary of the protrusion and base of the ruffle. Intensity measurements for ruffles were carried out by manually defining regions of interest (ROI) that consisted of PA-Rac1 signal, and the intensity of second channel pertaining to different myosin types was extracted. A similarly sized ROI (equivalent in pixel numbers) was chosen as a cytosol control to normalize. Intensities were plotted in Origin Pro. Cross-correlation between the signals in the two channels was performed using Origin Pro.

## RESULTS

### Continuous activation of Rac1 activity is required for effective ruffle formation

To visualise dorsal membrane ruffling on-demand, we imaged RPE1 cells transiently transfected with p TriExmCherry PA-Rac1 as previously described (15, 16). This enabled us to optogenetically activate Rac1 signalling and induce dorsal ruffling of the membrane as well as allowing us to visualise this process using fluorescence imaging. To visualise the dynamic and transient Rac1 ruffles on the dorsal surface of live cells, we resorted to lattice light sheet microscopy (26), which provides rapid and photo-gentle volumetric imaging at near diffraction limited resolution (Fig. 1A). To produce precise temporal activation of PA-Rac1, we attempted two strategies of optogenetic excitation. We attempted to activate PA-Rac1 using a 445 nm laser passing through the spatial light modulator and excitation objective to produce a ‘full length’ or ‘quarter length’ multi-bessel beam excitation profile (Fig. 1B, C). While effective photoactivation could be achieved in specific regions of the cell using other constructs (supplementary material), we did not observe any induced ruffling processes (Fig. 1D). We attributed this to the fact that using the SLM based photoactivation is discontinuous and only activated the molecules in imaging plane at that instance while scanning the entire volume. This resulted in insufficient local concentration of activated PA-Rac1 to generate ruffles. Although the dark-state recovery kinetics of As Lov2 are over a minute, the threshold local concentration for sheet-like ruffles to be organized is never achieved presumably owing to the requirements of downstream interactions, cytoskeletal events including polymerization leading to protrusion formation (Fig. 1C). A continuous excitation through the imaging sequence should allow sufficient time for enough number of molecules as well as subsequent downstream events of actin meshwork formation resulting in protrusion formation to take place. We, therefore, resorted to a wide-field excitation strategy focusing the 445 nm laser at the back focal plane of the detection objective (Fig. 1E, Methods). This produced continuous optogenetic stimulation that was independent of the volumetric imaging of the sample (Fig. 1E). This mode of photoactivation leads to effective PA-Rac1 activation and transformation of the dorsal surface of the cell from filopodial protrusions and small membrane ruffles to large-scale sheet-like membrane ruffles as has been previously reported (Fig. 1F, Supplementary movie 1) (15, 16). All experiments described below were carried out using wide field (continuous) photoactivation.

**Figure 1.**
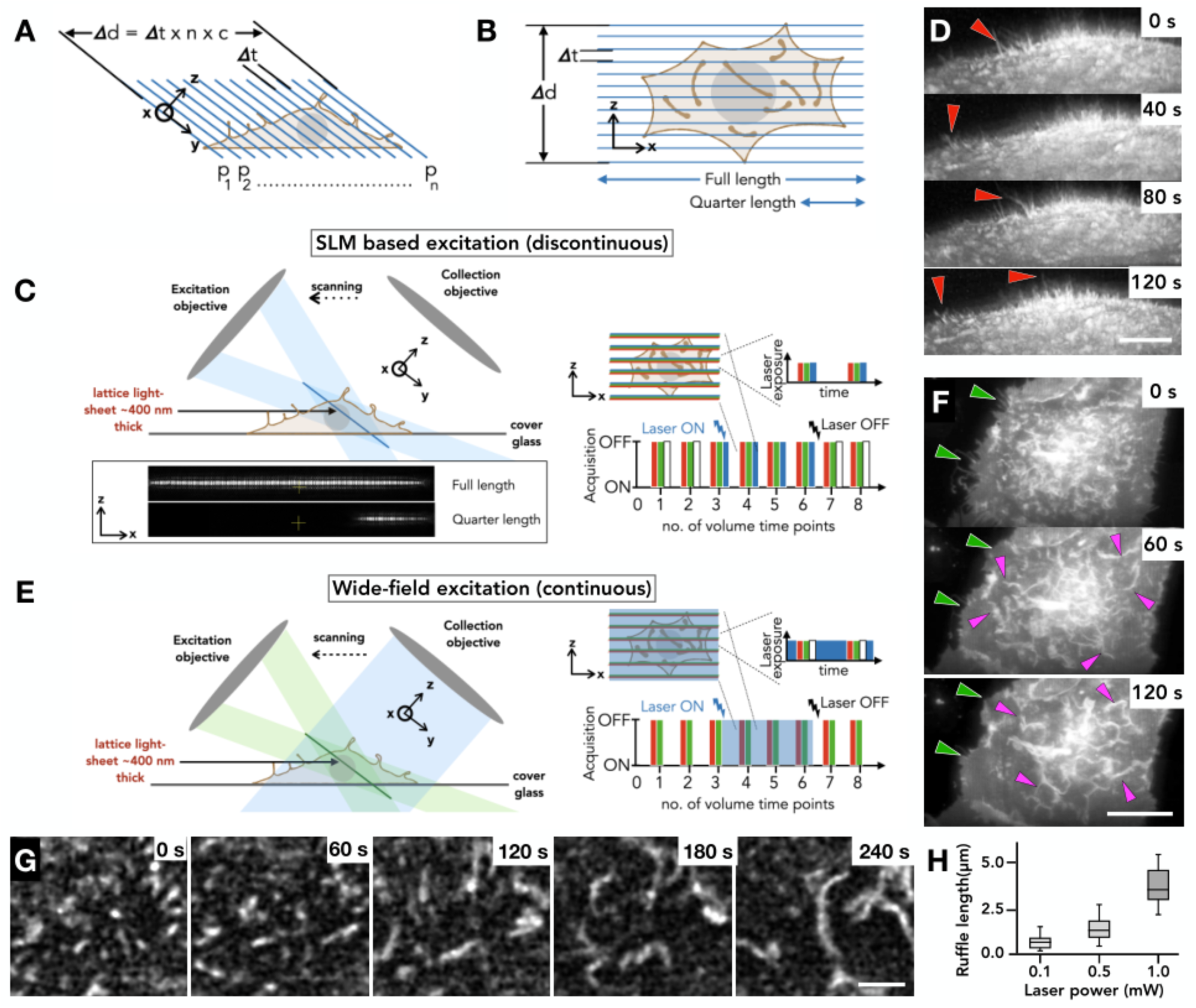
Optogenetic stimulation of PA-Rac1 on a Lattice Light Sheet Microscope. ***A***. Volumetric acquisition using LLSM. ***B***. Top-down view of LLSM acquisition of a cell, in comparison to the lattice light sheet excitation plane. ***C***. Schematic representation of optogenetic excitation using the SLM to produce controlled 445 nm excitation, shown in right panel. Using the SLM in the excitation path results in discontinuous excitation ***D***. RPE1 cell stimulated using SLM-based excitation ((as described in (C) of PA-Rac1 does not suppress filopodia and no ruffles are observed. Scale bar = 5 µm ***E***. Schematic representation of widefield optogenetic excitation passed through the detection objective leading to continuous excitation, shown in right panel. ***F***. Dorsal membrane ruffles produced following wide field optogenetic stimulation (as described in E). Scale bar = 10 µm ***G***. Growth of PA-Rac1 ruffles in response to photoactivation. Scale bar = 3 µm. ***H***. Boxplot of PA-Rac1 ruffle length (µm) vs photoactivation laser power (mW). Outer range shows total range, box height shows the standard deviation, and inner bar shows mean.

Efficient ruffle formation involves the following process: (a) there is an overall decrease in the filopodia, and punctate structures appear that grow along its length (Fig. 1G). (b) At this regime, the characteristic length of the ruffles is dependent on laser power (Fig. 1H). (c) At higher laser powers, the ruffles grow and merge with one another, resulting in formation of complex branched geometries (Supplementary movie 2). PA-Rac1 was found to be membrane localized, as previously reported [14], and enriched in the ruffles following photoactivation (Fig. 1F, 1G, Supplementary movie 1, 2) and as such PA-Rac1 mCherry signals can be used to visualize and identify the ruffles.

### Repetitive photoactivation sequences reveal re-initiation of growth on pre-formed ruffles

Our observations suggest that with continuous excitation, a threshold concentration of activated PA Rac1 molecules is achieved that result in local cytoskeletal rearrangements. PA-Rac1 mediated actin reorganizations then lead to efficient lamella formation. This activated Rac1 dependent process represents a barrier that cannot be overcome in discontinuous photoactivations. We reasoned that a ruffle generated in the first cycle would have molecular arrangements and actin meshwork architecture that would be pre-disposed for re-initiation of a collapsing ruffle. To test this, we performed repeated consecutive stimulation experiments on cells expressing PA-Rac1 (Fig. 2A, B). Using VR, we quantified the length of distinguishable membrane protrusions (Fig. 2E, Supplementary movie 3). The first two stimulations did not lead to a discernible increase in the height of the ruffle protrusions, but rather an increase in the length that reflects an increase in net polymerization. The second and third stimulations led to a stepwise increase in the overall height of the membrane protrusions and extrusion of very long sheets on the dorsal surface. This suggests that pre-activated, but collapsing ruffles can be reactivated, leading to re-initiation of assembly of actin meshwork. The increase in the height of the protrusions support the idea of polymerization forces at play along the direction of protrusion when PA-Rac1 is activated. Further, upon turning off the activation laser after the third stimulation, the complex membrane folds collapsed onto very bright compact structures owing to actomyosin contractility (Fig. 2A, B, C, red boxes, Supplementary movie 4). Interestingly, following this collapse, activating the PA-Rac1 resulted in ‘unfurling’ of compact contracted membrane folds (Fig. 2C, stimulation 4, magenta and green arrows). These unfurling events showed rotational motions driven by the polymerization forces (Fig. 2C, D magenta and green arrows, Supplementary movie 5). These observations clearly demonstrate that PA-Rac1 activation results in a polymerization dominant phase that is capable of stretching the membrane resulting in unfolding of complex membrane folds.

**Figure 2.**
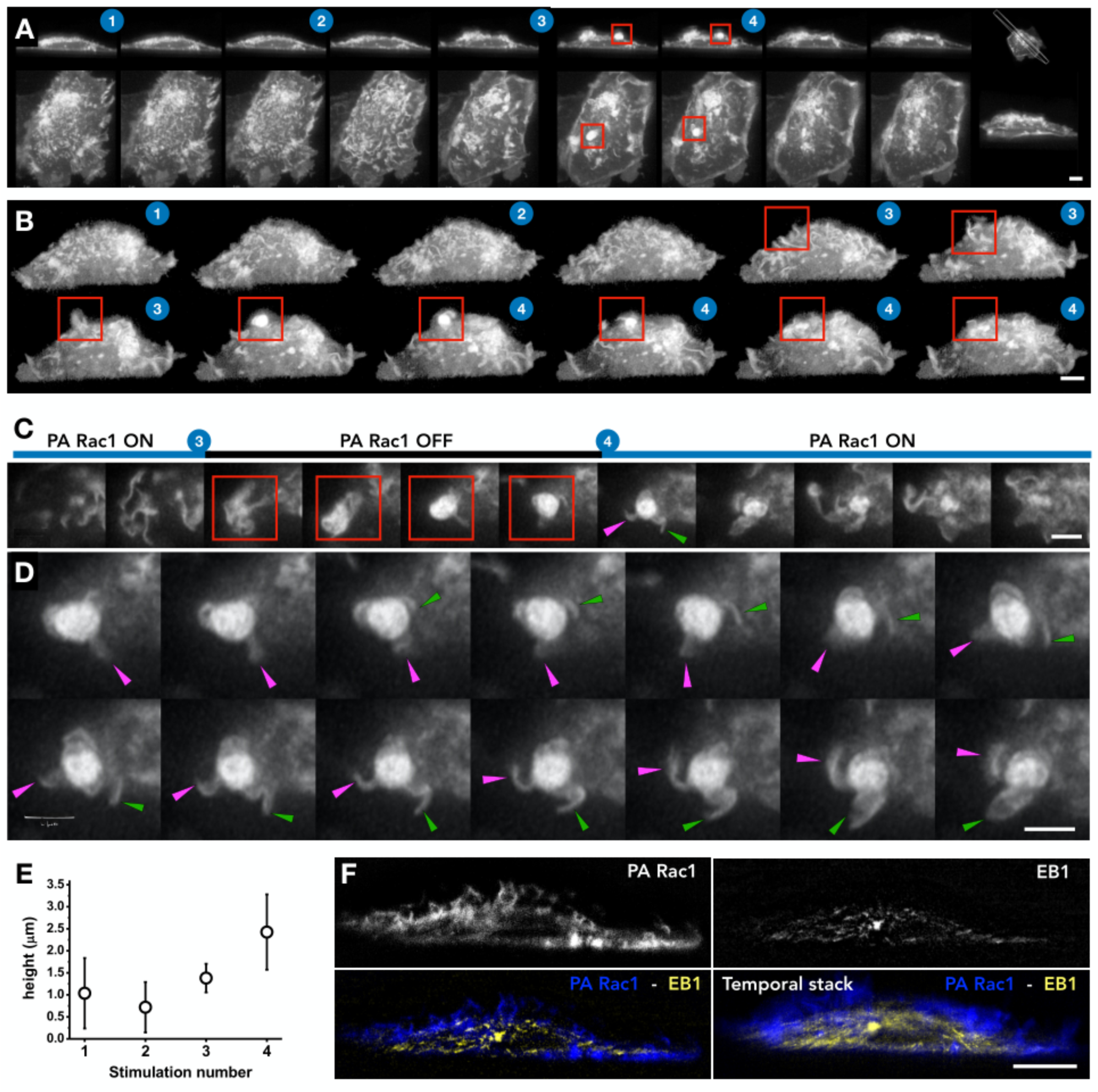
Repeated optogenetic activation leads to incremental increases in protrusion length. ***A***. Montage of PA-Rac1 membrane ruffles in whole RPE1 cells in response to multiple optogenetic stimulations (supplementary movie 3). Left to right shows changes in Rac1 with time. Top panel shows a 2D slice through the cell (as marked in the last panel). Bottom panels shows Top views. The blue number overlays indicate the sequential optogenetic stimulation. The red square highlights a collapse following a photoactivation laser switch off. Scale bars for both panels = 5 µm. ***B***. Montage of ruffle evolution in response to multiple optogenetic stimulations of PA-Rac1. With multiple excitations, the ruffles become longer, extruding larger sheet like protrusions. Scale bar = 5 µm. ***C***. A temporal zoomed montage of red box in A-C. Formation of highly complex collapsed membrane folds post PA-Rac1 inactivation (3) and unfurling of the complex folds upon re-activation of PA-Rac1 (4). Frames every 20 s ***D***. Unfurling of complex membrane folds undergoing rotational movements with polymerization at ‘leading edges’ of the ruffles. The magenta and green arrows follow the same protrusions across the montage. Frames every 4 s. ***E***. Graph of ruffle height vs the number of times stimulated with 445 nm laser. Error bars indicate standard deviation. ***F***. Cross-sectional slice of RPE1 cell transfected with PA-Rac1 (blue) and EB1 (yellow). Bottom right panel shows a temporal stack MIP over 10 min. scale bar = 5 µm.

While lamellipodial protrusions are mainly actin driven structures, in some cases also a microtubule association was also observed (27, 28). To investigate whether dynamic microtubules are associated with the PA-Rac1 membrane protrusions, we co-transfected cells with EB1-GFP and investigated the localisation and dynamics of protrusions in response to the optogenetic activation of PA-Rac1. We observed no microtubules entering the dorsal ruffles (Fig. 2F, Supplementary movie 6). The absence of microtubule-ruffle interactions may be attributed to the transient nature of the ruffles, the sharp angle of propagation between the microtubules and ruffles or the dense actin meshwork architecture preventing microtubule entry. These observations taken together suggest that actin machinery is the dominant structural component orchestrating membrane ruffle dynamics.

### PA-Rac1 produces larger and straighter membrane protrusions than PMA

PA-Rac1 induced ruffle dynamics strongly suggest that actin polymerization forces dominate during the PA-Rac1 activated phase. To characterize the mechanical properties and the dimensions of the membrane ruffles, We compared ruffles that are stimulated using photoactivation of PA-Rac1 with those formed by stimulation with 100 nM PMA, a chemical agonist that induces membrane ruffling (29) (Fig. 3A). Upon Rac1 activation, we observed templated assembly of dorsal ruffles which polymerised from lamellipodia at the cell periphery before ‘lifting off’ and propagating along the dorsal cell surface (Fig. 3C, Supplementary movie 7) as also observed by Begemann et al. (30). Moreover, we also noted that dynamic membrane protrusions formed across the dorsal surface in response to PA-Rac1 activation (Fig. 3B). The observed membrane transformations are complex and developing image analysis-based algorithms to precisely segment out such dynamic pleomorphic dynamic structures with varying fluorescence intensities present various difficulties. We resorted to a Virtual Reality (VR) based approach (24), where the three-dimensional movies were rendered to enable the depth perception in the volumetric images akin to naturally evolved depth perception in vision of our surroundings. Using VR, we measured the end-to-end distances and the contour lengths of the ruffles as viewed from the top (Fig. 3D). Although the membrane ruffle is a sheet-like structure that is attached to the basal substrate in a very complex manner through acto-myosin cortex and other cytoskeletal structures, its two-dimensional conformation in the membrane plane is strikingly similar to that of a semiflexible filament (Fig. 3E). This allowed us to quantify their 2D conformations of different activations in terms of the persistence length ***l***_***p***_, by analysing their squared end-to-end distance 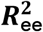 as a function of the contour length ***L*** using the expression for the 2D worm-like chain model (Fig. 3F) (25, 31): 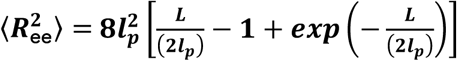.

**Figure 3.**
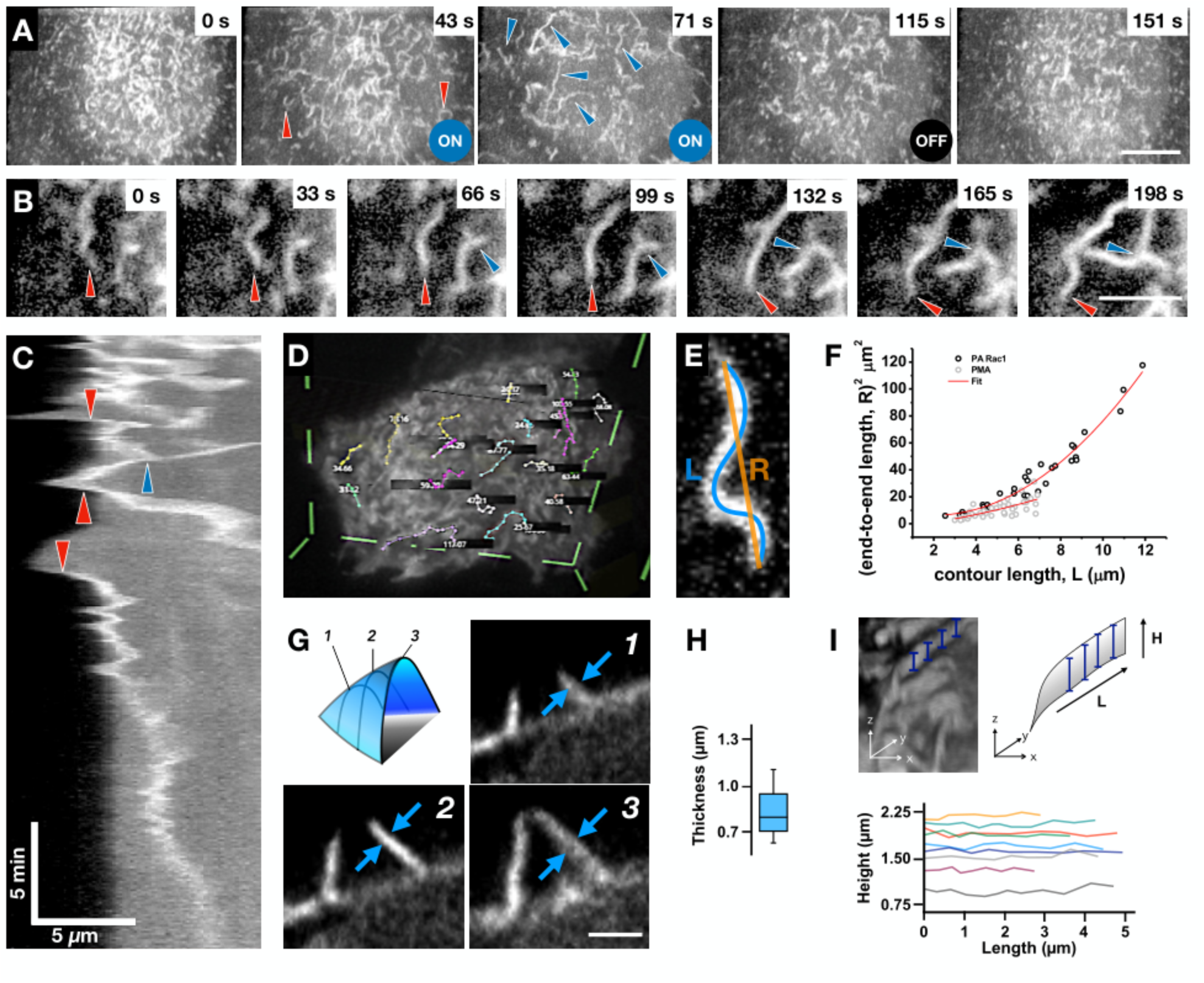
PA-Rac1 ruffles are lamellar structures that are formed and maintained by actin polymerization. **A**. Representative montage of ruffle formation and elongation in response to optogenetic stimulation in RPE1 cells. Red arrows indicate highly curved membrane ruffles stimulated by PMA; blue arrows indicate large ruffles formed in response to optogenetic stimulation. Scale Bar = 10 µm. ***B***. Representative montage of PA-Rac1 stimulated ruffle dynamics. Red arrows show the growing end of the ruffle, blue arrows show branching points. Scale bar = 3 µm. ***C***. Kymograph of PA-Rac1 signal at a lamellipodial edge during Rac1 stimulation. The Red arrows represent Lift-Offs of lamellipodia (LOL) and blue arrow points at a merger of two ruffles. X scale bar = 5 µm, Y scale bar = 5 min. ***D***. Representative screen capture showing measurements of membrane ruffle contours using Syglass VR. ***E***. Representative diagram showing squared end-to-end length (R), and contour length (L) overlaid on a top view of a ruffle. ***F***. Squared end-to-end length vs. Contour length of ruffles stimulated by PMA (grey) or PA-Rac1 (black). Curve of fit (red), corresponds to the persistence length derived from the 2D worm-like chain model. ***G***. Representative images showing the measurement of ruffle thickness (blue arrows). Image panels show slices through a macropinocytic cup as portrayed in the top left panel. Scale bar = 3 µm. ***H***. Boxplot of membrane ruffle thickness (µm). Outer range shows total range, box height shows the standard deviation, and inner bar shows mean. ***I***. Representative images showing extraction of ruffle protrusion height. Measured height shown by blue bars top right, overlay top left. Bottom shows ruffle height (µm) vs length (µm). Measurements were taken for 9 representative ruffles.

The persistence length of PA-Rac1 ruffles (∼5 µm) is considerably larger than that of PMA-activated ruffles (∼0.8 µm) (Fig. 3F), which perfectly agrees with their visual perception in Figs. 2A. At the same time this does not necessarily means that PA-Rac1 ruffles are correspondingly stiffer: the PMA ruffles may be equally stiff but have a smaller persistence length owing to their intrinsic curvature, potentially due to concomitant myosin activity. Then the observed persistence length ***l***_p,obs_ is related to the persistence lengths corresponding to the bending stiffness of the structure and its intrinsic curvature, ***l***_***p***_ and ***l***_p,int_, respectively, via the relationship 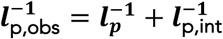 (32, 33). Assuming that the PMA ruffles have ***l***_***p***_ close to that of the PA-Rac1 ruffles, we can estimate the intrinsic curvature-related persistence length of PMA ruffles as ∼1 µm.

We also measured the height of the ruffle along the length as well as the thickness in deconvolved images (Fig. 3H,I). We observed that whilst membrane ruffles showed variation in their height during the encapsulation process, they showed a rather uniform and consistent thickness with a mean of 0.83 μm across their length (Fig. 3G,H). The PA-Rac1−based dorsal ruffles showed a characteristic width similar to that observed in neutrophil-like HL-60 cells (0.78 μm by LLSM, 0.54 μm by spinning-disk and 0.64 μm by SEM) (18, 34), suggesting that there exists a linear arrangement of regulatory molecules that produces ruffles in a similar manner. The PA-Rac1 activation may drive an increase in polymerization leading to the slightly thicker sheets observed in our experiments. Further, the variation in height over few micrometers of ruffle length was below 110 nm, as measured in deconvolved data sets using segmentations based on thresholding that can delineate edges with sub-pixel resolution (Fig. 3I, supplementary movie 8). This suggests that the PA-Rac1−generated ruffles are devoid of any tent-pole like arrangements (35) and can sustain dependence on polymerization forces. The relatively small standard deviation of the height along the length of the ruffles can be attributed to complex interplay between actin polymerization at the edge of the ruffle and the linear sheet-shape of the ruffle. Since Rac1 activity is known to increase Arp2/3-based branched actin meshwork (36, 37), a noise suppression system involving these parameters may be envisaged (38).

Our observations of intrinsically sheet-like ruffles of several micrometres long, generated by PA-Rac1 suggests that protrusion formation can also take place with initiation coming from chemical activation of Rac1 than mechanical curvature dependent activation alone (30). In this case, we have used PA-Rac1, but across the diverse roles of Rac1-driven ruffling, there exist multiple mechanisms of activation to produce migration (39, 40), micropinocytosis (3) or macrophage tunnelling nanotube biogenesis (41).

There is no consensus on what distinguishing factors determine circular and linear dorsal ruffles apart from their geometry. Linear ruffles are generally associated with the lamellipodia and circular dorsal ruffles are typically stimulated by an agonist (5). We consistently observed that activated PA-Rac1 resulted in stiffer and more linear ruffles, whereas switching off PA-Rac1 caused ruffles to fold over, showing rotational features characteristic of circular dorsal ruffles (42). This suggests that the rotational characteristics that define ‘circular’ dorsal ruffles may be a complex interplay of polymerization and contractile forces.

### Deactivation of PA-Rac1 leads to buckling and closure

We hypothesized that the reason for the ruffles formed by PMA to be consistently curved as compared to PA-Rac1−induced ruffles may be the contractile forces that cause membrane sheet protrusions to fold and collapse leading to engulfment. Therefore, we tested whether increased polymerization-induced forces keep the ruffle in a ‘stretched’ geometry and therefore whether upon releasing the PA-Rac1 from the activated state, the contractile forces dominate which leads to ruffle collapse. To assist resolvable observations, we combined PMA and PA-Rac1−mediated stimulation, as above (Fig. 4A, Supplementary movie 7). During optogenetic PA-Rac1 activation, we observed that large dorsal ruffles formed at the cell periphery (Fig. 4A, white box 3B, white arrows). We observed that deactivating PA-Rac1 led to various behaviours suggestive of contractile forces, which included buckling (Fig. 4A,C, red arrows 3F, white arrows) and formation of cup-like structures (Fig. 4A, D, Magenta arrows). Further, upon switching off the photoactivation laser, splitting of ruffles (Fig. 4A, blue arrows) and subsequent recoil of the split ruffle was also observed that led to buckling and folding of the ruffle onto itself (Fig. 4A, E, green arrows). This led to cup closure and macropinosome formation (Fig. 4G, Supplementary movie 7,8,9)

**Figure 4.**
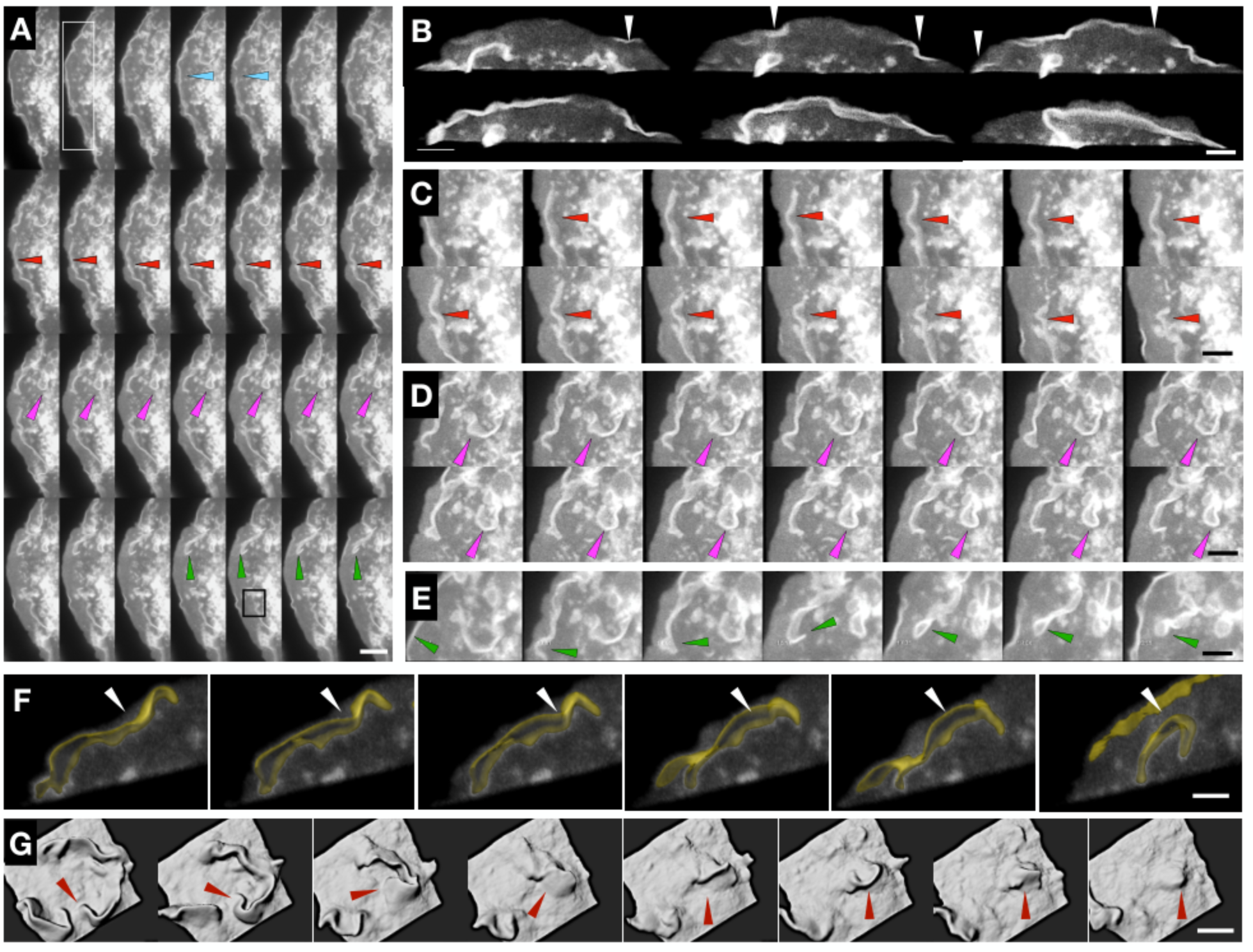
PMA and PA-Rac1 together give rise to complex coalescence of membrane protrusions. ***A***. Montage of PA-Rac1 ruffle coalescence in RPE1 cells treated with PMA (100 nM), figure runs from left to right, top to bottom (supplementary movie 7). Scale bar = 15 µm ***B***. Insert from Figure 3a. Marked by the white rectangle, montage of PA-Rac1 ruffle formation with optogenetic stimulation. Arrows indicate ruffle formation. Scale bar = 4 µm. ***C***. Insert from Figure 3a. Indicated by red arrows, montage of ruffle contraction post Rac1 stimulation. Scale bar= 5 µm. ***D***. Insert from Figure 3a. Marked by magenta arrows; montage of ruffle closing, indicated by magenta arrows. Scale bar = 5 µm. ***E***. Insert from Figure 3a, marked by the green arrows; montage of ruffle collapses following Rac1 inactivation, indicated by green arrows. Scale bar = 5 µm. ***F***. Insert from Figure 3a. Marked by black square; montage of ruffle buckling. The segmented ruffles are represented by yellow surface. Scale bar = 3 µm. ***G***. Montage of Rac1 ruffle collapse leading to cup-closure as a surface rendered volume. Scale bar = 4 µm.

Macropinocytosis, phagocytosis and ruffling are all processes that involve a substantial extension of sheet-like protrusions of an actin network-filled membrane that undergoes folding or wrapping to result in engulfment (1). At both mechanical and biochemical levels, these processes have a lot in common, including involvement of Rac1(2). Previous studies have shown that activation of Rac1 is sufficient to form macropinocytic or phagocytic cups, but deactivation or decline in Rac1 activity is necessary to lead to cup closure (16). In this regard, it is evident that the spatial coordination of cup closure is a significant task as it needs to occur at the right place without pushing out the engulfed fluid in macropinosome. In naturally occurring macropinocytosis, Rac1 deactivation may occur through coordinated recruitment of Rho GEFs and GAPs (43, 44). Here, using PA-Rac1, we show that the process of ruffling that leads to engulfment can be ‘held’ at the polymerization force dominated stage and can be released by switching off the photoactivation laser, resulting in contractile forces to dominate and cause engulfment.

### Myo1E associates with membrane ruffles with high spatio-temporal coincidence

To test if we can measure the partitioning of proteins specifically into the ruffles and their dynamics with respect to the formation of ruffles, we investigated the dynamic localisation of various myosins during the PA-Rac1−stimulated ruffling process. MyoIIA is not known to be part of ruffling machinery; however, both Myo1e and MyoX have been reported to localise to the phagocytic cup (45, 46). Both Myo1e and MyoX have PH-domain that can enable binding to PI(3,4,5)P3. Though phagocytic cups and macropinocytic cups are structurally similar — consisting of lamellar structures — we investigated the associations of particular myosins with PA-Rac1−based ruffles to ask whether there are distinguishing features. Therefore, we assayed if MyoX and Myo1e were specifically enriched in PA-Rac1 ruffles. We expressed MyoIIA, MyoX or Myo1e along with PA-Rac1, photoactivated PA-Rac1 (WF mode) and performed 2-colour imaging (Fig. 5D, Supplementary movie 10). We did not detect any enrichment of MyoIIA or MyoX, in comparison to Myo1e, which showed a high level of specific enrichment, similar to the PA-Rac1 YFP positive control (Fig. 5A). Myo1e localised to PA-Rac1 ruffles at the same time as its formation (Fig. 5 B, E). Correlation analysis showed a single peak at 0 s, indicating simultaneous recruitment of Myo1e to PA-Rac1 signal at time scales longer than the time-resolution of our measurements (8s) (Fig 5C). MyoX, on the other hand inhibited much of the independent ruffling processes with the measured ruffles originating from lamellipodia (data not shown).

**Figure 5.**
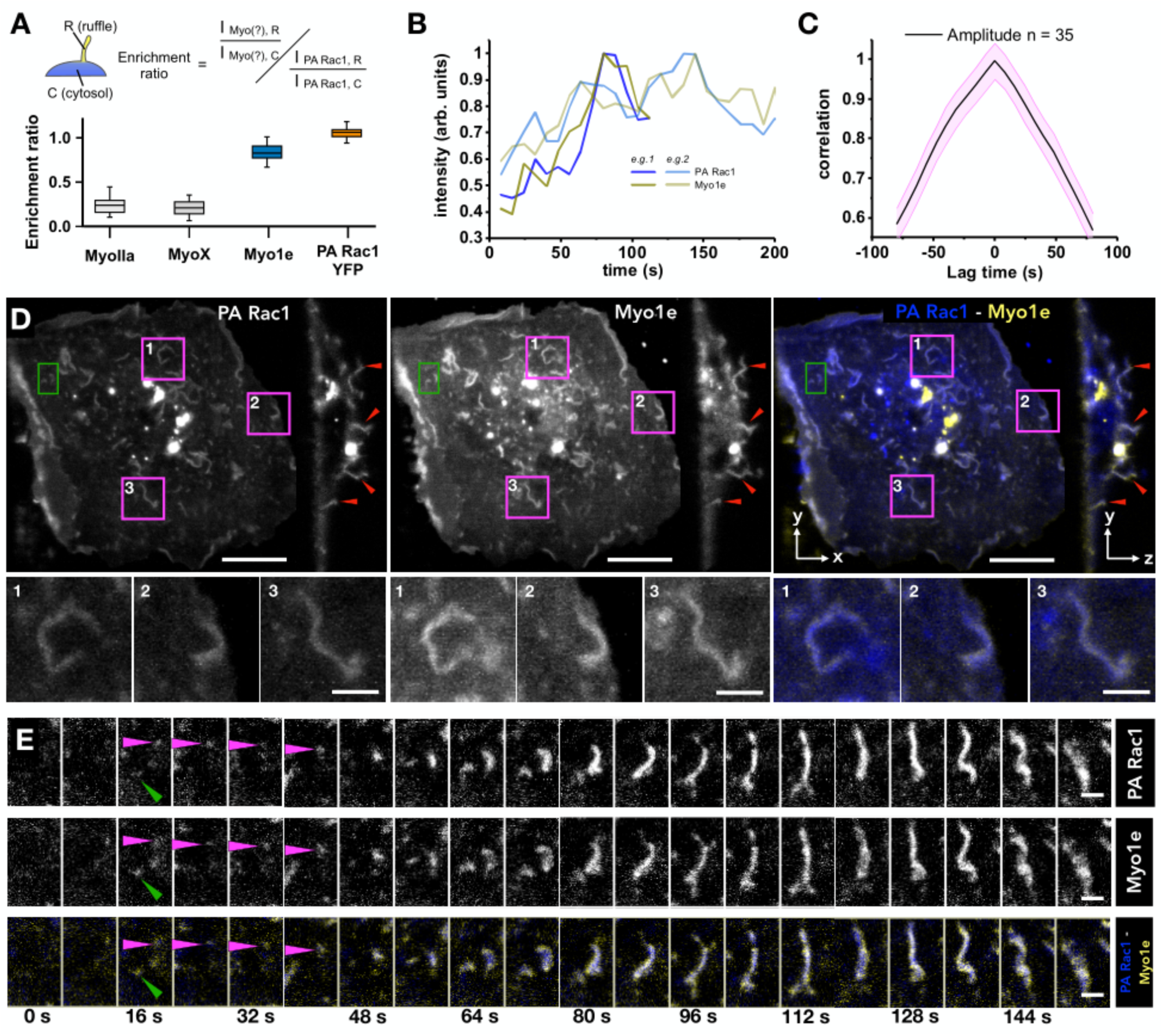
Myo1e colocalises to dorsal membrane ruffles. ***A***. Boxplot of the enrichment ratio of investigated proteins in membrane ruffles, as determined by the equation above (Methods). Bars show range, the inner box shows S.D, inner bar shows the mean value. n = 4 cells each. ***B***. Normalised PA-Rac1 and Myo1e intensities with time (seconds) for two example membrane ruffles. ***C***. Correlation of PA-Rac1 and Myo1e signals. Correlation measured from 35 ruffles ***D***. Top, MIP and cross-section of PA-Rac1 (blue) and Myo1e (yellow). Pink boxes refer to zoomed panels, bottom. Red arrows highlight colocalising dorsal ruffles. Green box refers to zoomed time series (Figure E). Scale bar = 5 µm ***E***. Time series of PA-Rac1 (blue) and Myo1e (yellow). Scale bar = 1.25 µm

**Figure 6.**
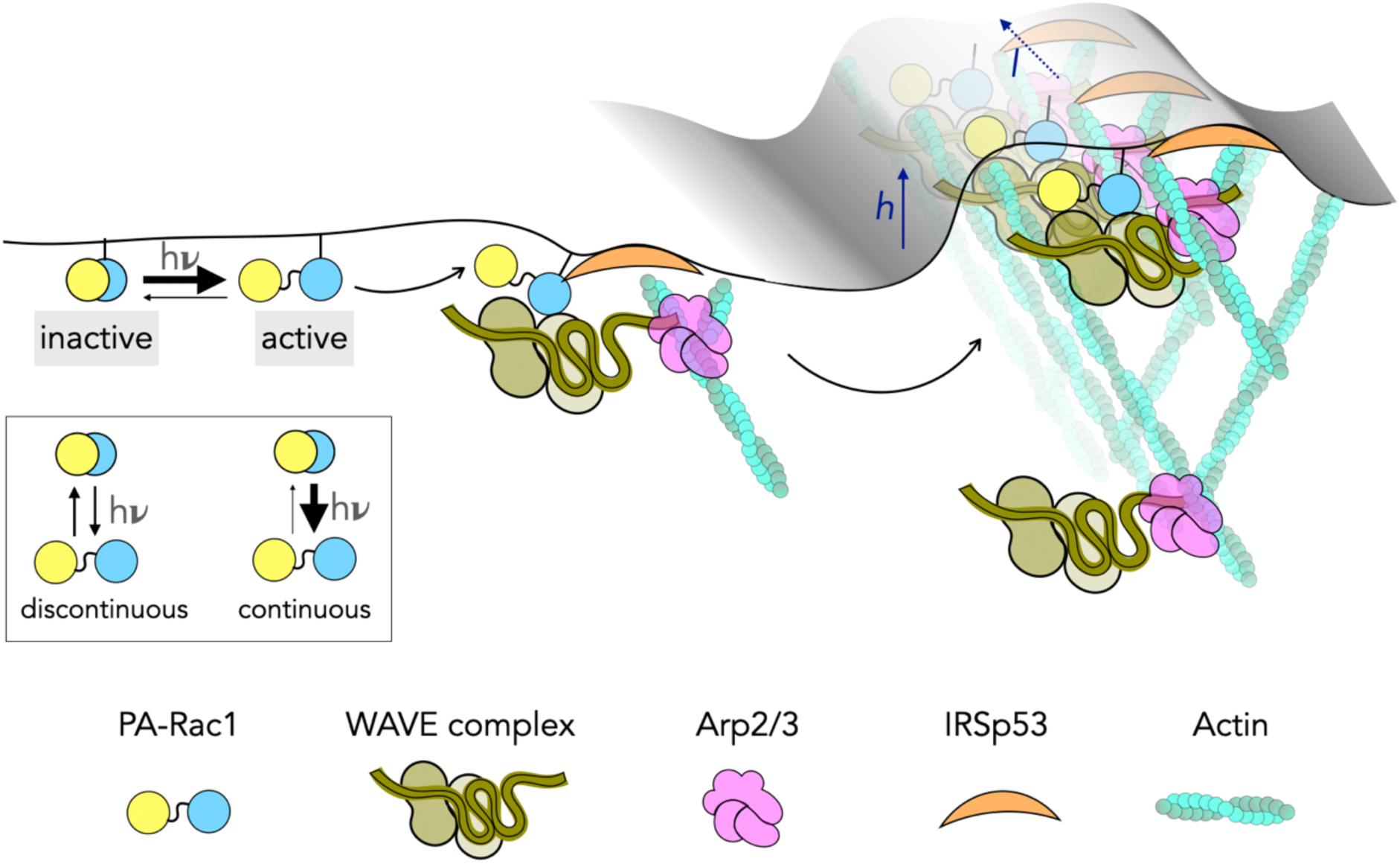
Summary figure of threshold-based assembly of lamellar ruffles generated by PA-Rac1.

Whilst MyoX has been reported to be localising in phagocytic cups, we could not detect any MyoX signal in PA-Rac1 based ruffles. However, the appearance of Myo1e in the ruffles is in agreement with a recent study that showed that Myo1e played a role in macropinocytic cup closure in the context of Fc receptor induced micropinocytosis (46). It is also of note that Myo1e or Myo1f have been implicated in lamellipodial neutrophil migration (47) and FAK-PI3K-RAC1 associated B-cell migration(48). Together, our results suggest that some general architectural components of the lamellar protrusion machinery may be conserved, and there may be other differences between phagocytic, pinocytic and lamellipodial protrusions with distinct sets of myosins engaged in different processes.

## CONCLUSION

We have combined LLSM with optogenetic activation for measuring the morphodynamics of PA-Rac1−based ruffling processes with high spatio-temporal resolution. Processes on the dorsal side of the cell are difficult to measure by many traditional geometries of imaging. We demonstrate that optogenetic approaches, where a process may need a threshold number of molecules (e.g. membrane ruffling) to be activated, can be achieved using simpler optical arrangements of wide field photoactivation, even though use of SLMs offer precision control over space and time. We also use VR-based analysis tools that we find to be extremely effective in manual measurements of complex morphologies in the three-dimensional volumes.

The results presented here clearly demonstrate that to achieve efficient ruffling, the photoactivation needs to be continuous. This is in line with previous studies that have shown that Rac1-GTP directly Interacts with WAVE with low affinity, and thus a high Rac1 concentration is required to activate the WAVE complex (36, 49, 50). From the results demonstrated in this study, along with previously reported molecular interactions that govern branched actin formation downstream of Rac1, a multi-step process can be envisaged. Photoactivated Rac1 triggers Arp2/3 mediated actin branching, which is presumably mediated through the WAVE regulatory complex (51, 52). With polymerized actin generating pushing forces against the membrane that can generate negative curvature, I-Bar protein IRSp53, which is known to interact with Rac1 and WAVE is recruited, which positively reinforces the protrusion geometry (30, 53, 54). A complex interplay between the membrane tension, polymerization forces of actin and organization of the branched actin meshwork through PA-Rac1 -IRSp53 – Wave – Arp2/3 is envisaged that can lead to generation of lamellar protrusion. This polymerization is sustained as long as PA-Rac1 is active. This organizational architecture is maintained at least for a minute even after switching off the photoactivation laser, which results in re-initiation of polymerization at the lamella edges.

Rac1 activation is critical in a number of distinct processes that involve transient actin-driven membrane protrusions including receptor-stimulated macropinocytosis, pinocytosis, lamellipodial driven migration (16, 39, 55). Photoactivation allows disentangling time and activity-dependent processes as demonstrated here by PA-Rac1. Our approach, demonstrated here, is widely applicable to investigate processes occurring at the dorsal surface of the cell, more accessible owing to the imaging geometries of the light-sheet based microscopes. We believe optogenetics, in combination with light-sheet approaches, can be instrumental in studying the spatio-temporal activity, localization and function of various other proteins, morphodynamics of protrusions and other complex phenomenology associated with the unsupported dorsal membrane surface.

## ABBREVIATIONS

CDR: Circular Dorsal Ruffle
LLSM: Lattice Light Sheet Microscopy
LOL: Lift-Offs of Lamellipodia
LOV: Light Oxygen voltage
PA-Rac1: Photoactivatable Rac1
PMA: Phorbol 12-myristate 13-acetate
ROI: Region of interest
SEM: Scanning Electron Microscopy
VR: Virtual Reality

## ACKNOWLEDGMENTS

HY is supported by an Australian Government Research Training (RTP) Scholarship. AP, UK are supported by Monash Biomedicine Discovery Institute Scholarships. The EMBL Australia Partnership Laboratory (EMBL Australia) is supported by the National Collaborative Research Infrastructure Strategy of the Australian Government.

## AUTHOR CONTRIBUTIONS

SA devised and directed the study, FL, SU, HY & UK performed LLSM experiments. HY, AP, HG, EPP, SA analysed data, SA, HY and EPP wrote the manuscript.

## SUPPLEMENTARY INFORMATION

**Supplementary Movie 1**. Transformation of the dorsal surface of the cells displaying filopodia to membrane ruffles upon photoactivation of PA-Rac1 expressed in RPE-1 cells.

**Supplementary Movie 2**. Photoactivating PA-Rac1 results in lamellipodial extension, generation and merging of ruffles.

**Supplementary Movie 3**. Repetitive photoactivation extrudes very long lamellar sheets.

**Supplementary Movie 4**. Sequential photo-activation cycles lead to long ruffles undergoing collapse into dense membrane structures that unfurl on re-initiation of ruffle growth by photoactivation.

**Supplementary Movie 5**. Polymerisation forces can drive rotational motions in unfurling complex membrane collapse formed as a result of sequences of photoactivation.

**Supplementary Movie 6**. EB1 GFP (Yellow) do not enter ruffles (Blue) generated by photoactivation.

**Supplementary Movie 7**. Ruffles simulated by photoactivation in cells expressing PA-Rac1 with 100 nM PMA in the media displaying lift-offs of lamellipodia and collapses leading to macropinosomes.

**Supplementary Movie 8**. Surface rendering of a ruffle following photoactivation undergoing collapse. Note the constant height of the ruffle.

**Supplementary Movie 9**. Sequences of on-off of photoactivation leading to a long ruffle undergoing a split, recoil and folding onto itself to form a versicle.

**Supplementary Movie10**. Myo1e (Yellow) is enriched in membrane ruffles (Magenta).

## Notes

### Competing Interest Statement

The authors have declared no competing interest.

### Summary of Updates

Title - grammar.

## REFERENCES

1. Swanson JA, Watts C. Macropinocytosis. Trends in cell biology. 1995;5(11):424–8.

2. Swanson JA. Shaping cups into phagosomes and macropinosomes. Nature reviews Molecular cell biology. 2008;9(8):639–49.

3. Yoshida S, Hoppe AD, Araki N, Swanson JA. Sequential signaling in plasma-membrane domains during macropinosome formation in macrophages. Journal of cell science. 2009;122(18):3250–61.

4. Ridley AJ. Membrane ruffling and signal transduction. Bioessays. 1994;16(5):321–7.

5. Buccione R, Orth JD, McNiven MA. Foot and mouth: podosomes, invadopodia and circular dorsal ruffles. Nature reviews Molecular cell biology. 2004;5(8):647–57.

6. Takenawa T, Suetsugu S. The WASP–WAVE protein network: connecting the membrane to the cytoskeleton. Nature reviews Molecular cell biology. 2007;8(1):37–48.

7. Hoon J-L, Wong W-K, Koh C-G. Functions and regulation of circular dorsal ruffles. Molecular and cellular biology. 2012;32(21):4246–57.

8. Ridley AJ, Paterson HF, Johnston CL, Diekmann D, Hall A. The small GTP-binding protein rac regulates growth factor-induced membrane ruffling. Cell. 1992;70(3):401–10.

9. Sun P, Yamamoto H, Suetsugu S, Miki H, Takenawa T, Endo T. Small GTPase Rah/Rab34 is associated with membrane ruffles and macropinosomes and promotes macropinosome formation. Journal of Biological Chemistry. 2003;278(6):4063–71.

10. Lanzetti L, Palamidessi A, Areces L, Scita G, Di Fiore PP. Rab5 is a signalling GTPase involved in actin remodelling by receptor tyrosine kinases. Nature. 2004;429(6989):309–14.

11. Ahram M, Sameni M, Qiu R-G, Linebaugh B, Kirn D, Sloane BF. Rac1-induced endocytosis is associated with intracellular proteolysis during migration through a three-dimensional matrix. Experimental cell research. 2000;260(2):292–303.

12. West MA, Prescott AR, Eskelinen E-L, Ridley AJ, Watts C. Rac is required for constitutive macropinocytosis by dendritic cells but does not control its downregulation. Current Biology. 2000;10(14):839–48.

13. Dharmawardhane S, Schurmann A, Sells MA, Chernoff J, Schmid SL, Bokoch GM. Regulation of macropinocytosis by p21-activated kinase-1. Molecular biology of the cell. 2000;11(10):3341–52.

14. Wu YI, Frey D, Lungu OI, Jaehrig A, Schlichting I, Kuhlman B, Hahn KM. A genetically encoded photoactivatable Rac controls the motility of living cells. Nature. 2009;461(7260):104–8.

15. Wu YI, Wang X, He L, Montell D, Hahn KM. Spatiotemporal control of small GTPases with light using the LOV domain. Methods in enzymology. 497: Elsevier; 2011. p. 393–407.

16. Fujii M, Kawai K, Egami Y, Araki N. Dissecting the roles of Rac1 activation and deactivation in macropinocytosis using microscopic photo-manipulation. Scientific reports. 2013;3:2385.

17. Ikeda Y, Kawai K, Ikawa A, Kawamoto K, Egami Y, Araki N. Rac1 switching at the right time and location is essential for Fcγ receptor-mediated phagosome formation. Journal of cell science. 2017;130(15):2530–40.

18. Fritz-Laylin LK, Riel-Mehan M, Chen B-C, Lord SJ, Goddard TD, Ferrin TE, Nicholson-Dykstra SM, Higgs H, Johnson GT, Betzig E. Actin-based protrusions of migrating neutrophils are intrinsically lamellar and facilitate direction changes. Elife. 2017;6:e26990.

19. Piehl M, Cassimeris L. Organization and dynamics of growing microtubule plus ends during early mitosis. Mol Biol Cell. 2003;14(3):916–25.

20. Taylor MJ, Perrais D, Merrifield CJ. A high precision survey of the molecular dynamics of mammalian clathrin-mediated endocytosis. PLoS Biol. 2011;9(3):e1000604.

21. Jacobelli J, Bennett FC, Pandurangi P, Tooley AJ, Krummel MF. Myosin-IIA and ICAM-1 regulate the interchange between two distinct modes of T cell migration. J Immunol. 2009;182(4):2041–50.

22. Bennett RD, Mauer AS, Strehler EE. Calmodulin-like protein increases filopodia-dependent cell motility via up-regulation of myosin-10. J Biol Chem. 2007;282(5):3205–12.

23. Arganda-Carreras I, Fernández-González R, Muñoz-Barrutia A, Ortiz-De-Solorzano C. 3D reconstruction of histological sections: Application to mammary gland tissue. Microscopy research and technique. 2010;73(11):1019–29.

24. Pidhorskyi S, Morehead M, Jones Q, Spirou G, Doretto G. syGlass: interactive exploration of multidimensional images using virtual reality Head-mounted displays. arXiv preprint 180408197. 2018.

25. Rivetti C, Guthold M, Bustamante C. Scanning force microscopy of DNA deposited onto mica: EquilibrationversusKinetic trapping studied by statistical polymer chain analysis. Journal of molecular biology. 1996;264(5):919–32.

26. Chen B-C, Legant WR, Wang K, Shao L, Milkie DE, Davidson MW, Janetopoulos C, Wu XS, Hammer JA, Liu Z. Lattice light-sheet microscopy: imaging molecules to embryos at high spatiotemporal resolution. Science. 2014;346(6208).

27. Waterman-Storer CM, Worthylake RA, Liu BP, Burridge K, Salmon E. Microtubule growth activates Rac1 to promote lamellipodial protrusion in fibroblasts. Nature cell biology. 1999;1(1):45–50.

28. Rinnerthaler G, Geiger B, Small J. Contact formation during fibroblast locomotion: involvement of membrane ruffles and microtubules. The Journal of cell biology. 1988;106(3):747–60.

29. Swanson JA. Phorbol esters stimulate macropinocytosis and solute flow through macrophages. Journal of cell science. 1989;94(1):135–42.

30. Begemann I, Saha T, Lamparter L, Rathmann I, Grill D, Golbach L, Rasch C, Keller U, Trappmann B, Matis M, Gerke V, Klingauf J, Galic M. Mechanochemical self-organization determines search pattern in migratory cells. Nature Physics. 2019;15(8):848–57.

31. Van der Maarel JR. Introduction to biopolymer physics: World Scientific Publishing Company; 2007.

32. Trifonov E, Tan R, Harvey S. Static persistence length of DNA. Structure and expression: proceedings of the Fifth Conversation in the Discipline Biomolecular Stereodynamics held at the State University of New York at Albany, June 2-6, 1987/edited by MH Sarma & RH Sarma. 1988.

33. Schellman JA, Harvey SC. Static contributions to the persistence length of DNA and dynamic contributions to DNA curvature. Biophysical chemistry. 1995;55(1-2):95–114.

34. Fleck RA, Romero-Steiner S, Nahm MH. Use of HL-60 Cell Line To Measure Opsonic Capacity of Pneumococcal Antibodies. Clinical and Diagnostic Laboratory Immunology. 2005;12(1):19.

35. Condon ND, Heddleston JM, Chew T-L, Luo L, McPherson PS, Ioannou MS, Hodgson L, Stow JL, Wall AA. Macropinosome formation by tent pole ruffling in macrophages. The Journal of Cell Biology. 2018;217(11):3873.

36. Eden S, Rohatgi R, Podtelejnikov AV, Mann M, Kirschner MW. Mechanism of regulation of WAVE1-induced actin nucleation by Rac1 and Nck. Nature. 2002;418(6899):790–3.

37. Chen B, Chou H-T, Brautigam CA, Xing W, Yang S, Henry L, Doolittle LK, Walz T, Rosen MK. Rac1 GTPase activates the WAVE regulatory complex through two distinct binding sites. Elife. 2017;6:e29795.

38. Garner RM, Theriot JA. Leading edge stability in motile cells is an emergent property of branched actin network growth. bioRxiv. 2020:2020.08.22.262907.

39. Kanazawa S, Fujiwara T, Matsuzaki S, Shingaki K, Taniguchi M, Miyata S, Tohyama M, Sakai Y, Yano K, Hosokawa K. bFGF regulates PI3-kinase-Rac1-JNK pathway and promotes fibroblast migration in wound healing. PloS one. 2010;5(8):e12228.

40. Das S, Yin T, Yang Q, Zhang J, Wu YI, Yu J. Single-molecule tracking of small GTPase Rac1 uncovers spatial regulation of membrane translocation and mechanism for polarized signaling. Proceedings of the National Academy of Sciences. 2015;112(3):E267–E76.

41. Hanna SJ, McCoy-Simandle K, Miskolci V, Guo P, Cammer M, Hodgson L, Cox D. The Role of Rho-GTPases and actin polymerization during Macrophage Tunneling Nanotube Biogenesis. Scientific Reports. 2017;7(1):8547.

42. Bernitt E, Koh CG, Gov N, Döbereiner H-G. Dynamics of actin waves on patterned substrates: a quantitative analysis of circular dorsal ruffles. PloS one. 2015;10(1):e0115857.

43. deBakker CD, Haney LB, Kinchen JM, Grimsley C, Lu M, Klingele D, Hsu P-K, Chou B-K, Cheng L-C, Blangy AJCb. Phagocytosis of apoptotic cells is regulated by a UNC-73/TRIO-MIG-2/RhoG signaling module and armadillo repeats of CED-12/ELMO. 2004;14(24):2208–16.

44. Hall AB, Gakidis MAM, Glogauer M, Wilsbacher JL, Gao S, Swat W, Brugge JS. Requirements for Vav guanine nucleotide exchange factors and Rho GTPases in FcγR-and complement-mediated phagocytosis. Immunity. 2006;24(3):305–16.

45. Cox D, Berg JS, Cammer M, Chinegwundoh JO, Dale BM, Cheney RE, Greenberg S. Myosin X is a downstream effector of PI(3)K during phagocytosis. Nature Cell Biology. 2002;4(7):469–77.

46. Barger SR, Reilly NS, Shutova MS, Li Q, Maiuri P, Heddleston JM, Mooseker MS, Flavell RA, Svitkina T, Oakes PW, Krendel M, Gauthier NC. Membrane-cytoskeletal crosstalk mediated by myosin-I regulates adhesion turnover during phagocytosis. Nature Communications. 2019;10(1):1249.

47. Krendel M, Kim SV, Willinger T, Wang T, Kashgarian M, Flavell RA, Mooseker MS. Disruption of Myosin 1e promotes podocyte injury. Journal of the American Society of Nephrology. 2009;20(1):86–94.

48. Girón-Pérez DA, Vadillo E, Schnoor M, Santos-Argumedo L. Myo1e modulates the recruitment of activated B cells to inguinal lymph nodes. Journal of Cell Science. 2020;133(5):jcs235275.

49. Ismail AM, Padrick SB, Chen B, Umetani J, Rosen MK. The WAVE regulatory complex is inhibited. Nature structural & molecular biology. 2009;16(5):561–3.

50. Chen Z, Borek D, Padrick SB, Gomez TS, Metlagel Z, Ismail AM, Umetani J, Billadeau DD, Otwinowski Z, Rosen MK. Structure and control of the actin regulatory WAVE complex. Nature. 2010;468(7323):533–8.

51. Ridley AJ. Life at the leading edge. Cell. 2011;145(7):1012–22.

52. Campellone KG, Welch MD. A nucleator arms race: cellular control of actin assembly. Nature reviews Molecular cell biology. 2010;11(4):237–51.

53. Miki H, Yamaguchi H, Suetsugu S, Takenawa T. IRSp53 is an essential intermediate between Rac and WAVE in the regulation of membrane ruffling. Nature. 2000;408(6813):732–5.

54. Lebensohn AM, Kirschner MW. Activation of the WAVE complex by coincident signals controls actin assembly. Molecular cell. 2009;36(3):512–24.

55. Buday L, Wunderlich L, Tamás P. The Nck family of adapter proteins: regulators of actin cytoskeleton. Cell Signal. 2002;14(9):723–31.

